# Facilitation by a hERG blocker is induced by pore opening and operates by pore reopening

**DOI:** 10.1101/2022.05.26.493575

**Authors:** Kazuharu Furutani, Ryotaro Kawano, Minami Ichiwara, Ryo Adachi, Colleen E. Clancy, Jon T. Sack, Satomi Kita

**Author notes:** Correspondence to: Kazuharu Furutani, Ph.D., Department of Pharmacology, Faculty of Pharmaceutical Sciences, Tokushima Bunri University 180 Nishihama-Boji, Yamashiro-cho, Tokushima, 770-8514 Japan. A preprint of this article has been deposited in bioRxiv.

## Abstract

A drug that blocks the cardiac myocyte voltage-gated K^+^ channels encoded by the human Ether-à-go-go-Related Gene (hERG) carries a potential risk of long QT syndrome and life-threatening cardiac arrhythmia, including *Torsade de Points*. Interestingly, certain hERG blockers can also facilitate hERG activation to increase hERG currents, which may reduce proarrhythmic potential. However, the molecular mechanism involved in the facilitation effect of hERG blockers remains unclear. The hallmark feature of the facilitation effect by hERG blockers is that a depolarizing preconditioning pulse shifts voltage-dependence of hERG activation to more negative voltages. Here we utilize a D540K hERG mutant to study the mechanism of the facilitation effect. D540K hERG is activated by not only depolarization but also hyperpolarization. This unusual gating property of the D540K hERG channel enables testing of hypotheses about the mechanism by which voltage induces facilitation of hERG by blockers. With D540K hERG, we find that nifekalant, a hERG blocker and Class III antiarrhythmic agent, facilitates not only current activation by depolarization but also current activation by hyperpolarization. Our results indicate that induction of facilitation is coupled to pore opening, not voltage *per se*. We propose that a gated-access mechanism is involved in the voltage-dependence of induction of facilitation. This study provides a molecular mechanism for modulation of hERG channels by nifekalant, a clinically important antiarrhythmic agent.

**Significance statement:** Nifekalant is a clinically important antiarrhythmic agent and a hERG blocker which can also facilitate voltage-dependent activation of hERG channels after a preconditioning pulse. Here we show that the mechanism of action of the preconditioning pulse is to open a conductance gate to enable drug access to a facilitation site. Moreover, we find that facilitation increases hERG currents by altering pore dynamics, rather than acting through voltage sensors.

## Introduction

The human ether-à-go-go related gene (hERG) encodes a voltage-gated potassium channel Kv11.1 subunit. hERG channels underlie the rapid component of the cardiac delayed-rectifier potassium current (*I*_Kr_) that mediates the rapid repolarization phase during the ventricular action potential (Sanguinetti et al., 1995; Sanguinetti and Tristani-Firouzi, 2006; Trudeau et al., 1995; Vandenberg et al., 2012). Drugs that decrease hERG current delay cardiac action potential repolarization and can result in long-QT syndrome, potentially leading to fatal arrhythmias such as *Torsade de Pointes* (TdP) (Kannankeril et al., 2010; Roden, 2000; Roden, 2008; Sanguinetti et al., 1995; Sanguinetti and Tristani-Firouzi, 2006; Surawicz, 1989; Vandenberg et al., 2012). hERG channels are unusually promiscuous targets for structurally diverse drugs and designing drugs that do not inhibit hERG channels is a major challenge (Sanguinetti and Tristani-Firouzi, 2006; Vandenberg et al., 2012). Curiously, certain hERG blockers exhibit secondary agonistic effects on hERG current, called “facilitation” that increases channel current at potentials close to the threshold for channel activation (Carmeliet, 1993; Furutani et al., 2019; Furutani et al., 2011; Hosaka et al., 2007; Jiang et al., 1999; Perry et al., 2010; Yamakawa et al., 2012). This facilitation is proposed to increase *I*_Kr_ in ventricular myocytes during the repolarization phase of action potentials and lower the risk that a hERG blocker will cause arrhythmia (Furutani et al., 2019).

Nifekalant, a Class III antiarrhythmic agent, blocks and facilitates hERG channels. Nifekalant facilitation of hERG channels shifts the conductance-voltage relation negative by ~20 mV (Furutani et al., 2019; Furutani et al., 2011; Hosaka et al., 2007). The hallmark feature of hERG facilitation by drugs is that this drug effect is induced by a depolarizing preconditioning pulse. However, how drugs affect the voltage-dependence of hERG activation and why preconditioning is required remains unclear. Here, we consider three hypotheses for the molecular origin of the voltage dependence of preconditioning for facilitation: 1) the charged moiety of nifekalant, 2) movement of hERG voltage sensors, or 3) channel opening.

We test these hypotheses with a hERG mutant that alters the coupling between voltage sensor movement and pore opening. Sanguinetti et al. characterized an aspartate 540 to lysine (D540K) mutant, located on the S4-S5 linker, that can be activated unusually by hyperpolarization, in addition to relatively normal activation in response to depolarization (Mitcheson et al., 2000; Sanguinetti and Xu, 1999). The mechanism of hyperpolarization-induced opening of D540K hERG has identified open channel block and trapping properties of hERG blockers (Kamiya et al., 2006; Mitcheson et al., 2000; Perry et al., 2004; Witchel et al., 2004). Critical residues for nifekalant block and facilitation are distributed in the pore and S6 helix (Hosaka et al., 2007; Kamiya et al., 2006), far from hERG residue 540.

We first show that hyperpolarization-activated channel openings of D540K hERG enable nifekalant to access the binding site for nifekalant inhibition. We then demonstrate that strong hyperpolarization of D540K hERG, but not wild-type (WT) channels, promotes nifekalant’s facilitation effect. These studies indicate that opening of the intracellular activation gate favors the binding of nifekalant to facilitate hERG channels.

## Materials and Methods

### Cells

A human embryonic kidney (HEK) 293 cell line stably expressing hERG (hERG-HEK) was kindly provided by Dr. Craig T. January and maintained in Dulbecco’s Modified Eagle Medium (DMEM; Fujifilm Wako Pure Chemical Corp., Osaka, Japan) supplemented with 10% fetal bovine serum (FBS; biowest, Nuaillé, France) and 400 µg/ml G418 (InvivoGen, CA, USA) as previously described (Zhou et al., 1998). An inducible D540K hERG HEK293 cell line (T-REx D540K hERG HEK) was generated using the Flp-In T-REx system (Thermo Fisher Scientific, MA, USA). The D540K hERG mutant was made site-directed mutagenesis from the wild-type (WT) (GeneBank accession No. U04270, WT hERG pCEP4), sequenced, and subcloned into the expression vector (pcDNA5/FRT/TO; Thermo Fisher Scientific). Cell lines were maintained in cell-culture treated polystyrene dishes at 37°C in a 5% CO_2_ atmosphere in growth media composed of DMEM media containing 10% FBS, and 1% penicillin–streptomycin solution (Fujifilm Wako Pure Chemical Corp., Osaka, Japan). Twenty-four hours before experiments, 1 μg/mL doxycycline HCl (Sigma-Aldrich, MO, USA) was added to media to induce D540K hERG channel expression.

### Electrophysiology

HEK cells used for electrophysiological study were adhered to the poly-L-lysine-coated (MW 30,000-70,000, Sigma-Aldrich; 0.1 mg/mL for 2 hr at 37 ºC) coverslips (CS-5R, Warner Instruments Corp., MA, USA) in 12-well plates and transferred to a small recording chamber mounted on the stage of an inverted microscope (IX71, Olympus Corp., Tokyo, Japan), and were continuously superfused with HEPES-buffered Tyrode solution containing (in mM) 137 NaCl, 4 KCl, 1.8 CaCl_2_, 1 MgCl_2_, 10 glucose, and 10 HEPES (pH 7.4 with NaOH). Membrane currents were recorded in a whole-cell configuration established using pipette suction (Hamill et al., 1981). The borosilicate micropipette had a resistance of 2–3 MΩ when filled with the internal pipette solution contained (in mM) 120 KCl, 5.374 CaCl_2_, 1.75 MgCl_2_, 10 EGTA, 10 HEPES (pH 7.2 with KOH). Bath and Internal solution osmolarity were adjusted to ~ 300 mOsm/kg with sucrose in some experiments. Liquid junction potential with this internal solution was calculated as less than −4 mV (Liquid Junction Potential Calculator, https://swharden.com/software/LJPcalc/app/), and the off-set was not corrected. Series resistance was typically under 5 MΩ. Series resistance compensation was used when needed to constrain voltage error to <10 mV. Leak compensation was not used. Whole-cell recordings were performed using an Axopatch 200B patch-clamp amplifier, Digidata 1322A interface and pClamp 9 software (Molecular Devices, Sunnyvale, CA, USA). The data were stored on a computer hard disk and analyzed using Clampfit 9 (Molecular Devices) and Igor Pro 7 and 9 (WaveMetrics, OR, USA). Experiments to characterize hERG facilitation were performed at room temperature (22-25°C).

Nifekalant was obtained from Cayman Chemical (Ann Arbor, MI).

### Monitoring current facilitation

The voltage protocols are indicated in each Figure as insets. For determining the voltage dependence of hERG activation and drug actions, the voltage protocols consisted of 20-second cycles containing two voltage steps from the holding potential of −80 mV. The first voltage step is a 4-second “test pulse” to a voltage between +80 mV and +60 mV with 10 mV increments to activate hERG channels. Afterward, the cell is returned to −60 mV for 2 seconds to record tail currents. To test the induction of the facilitation effect by nifekalant, the preconditioning pulse to +60 mV or −160 mV for 4-second was applied 20 seconds before each test pulse.

To show the I-V relationships, outward currents induced by a test pulse of −60 to +60 mV were normalized to the currents evoked by voltage steps to +10 mV without nifekalant. Inward currents induced by a test pulse of −160 mV to −60 mV were normalized to the currents evoked by voltage steps to −160 mV without nifekalant. To show the G-V relationships (hERG activation curves), depolarization-activated tail currents recorded at −60 mV were normalized to the currents evoked by voltage steps to +10 mV without nifekalant. Hyperpolarization-activated tail currents recorded at −60 mV were normalized to the currents evoked by voltage steps to −160 mV without nifekalant. Fitting was carried out using Igor Pro 9 (WaveMetrics) that employs nonlinear least squares curve fitting via the Levenberg-Marquardt algorithm. G–V relations were fit with a one power Boltzmann:

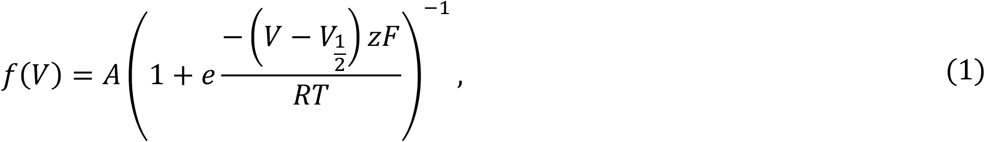

where *f*(*V*) is normalized conductance (*G*), A is maximum amplitude, *V*_1/2_ is activation midpoint, *z* is the valence in units of elementary charge (*e*_0_), *F* is the Faraday constant, *R* is the ideal gas constant, and *T* is absolute temperature.

The minimum Gibbs free energy (Δ*G*_Nif_) that nifekalant imparts to conductance, was calculated as

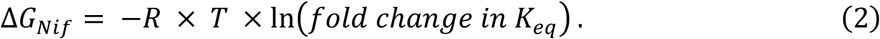

Here, *R* = 0.00199 kcal/(K·mol), and *T* = 298 K. *K*_eq_, or the equilibrium constant of channel opening, was approximated by the relative conductance of hERG either before or after induction of facilitation by nifekalant at *V*_1/2_ of block (Table 1).

**Table 1.**
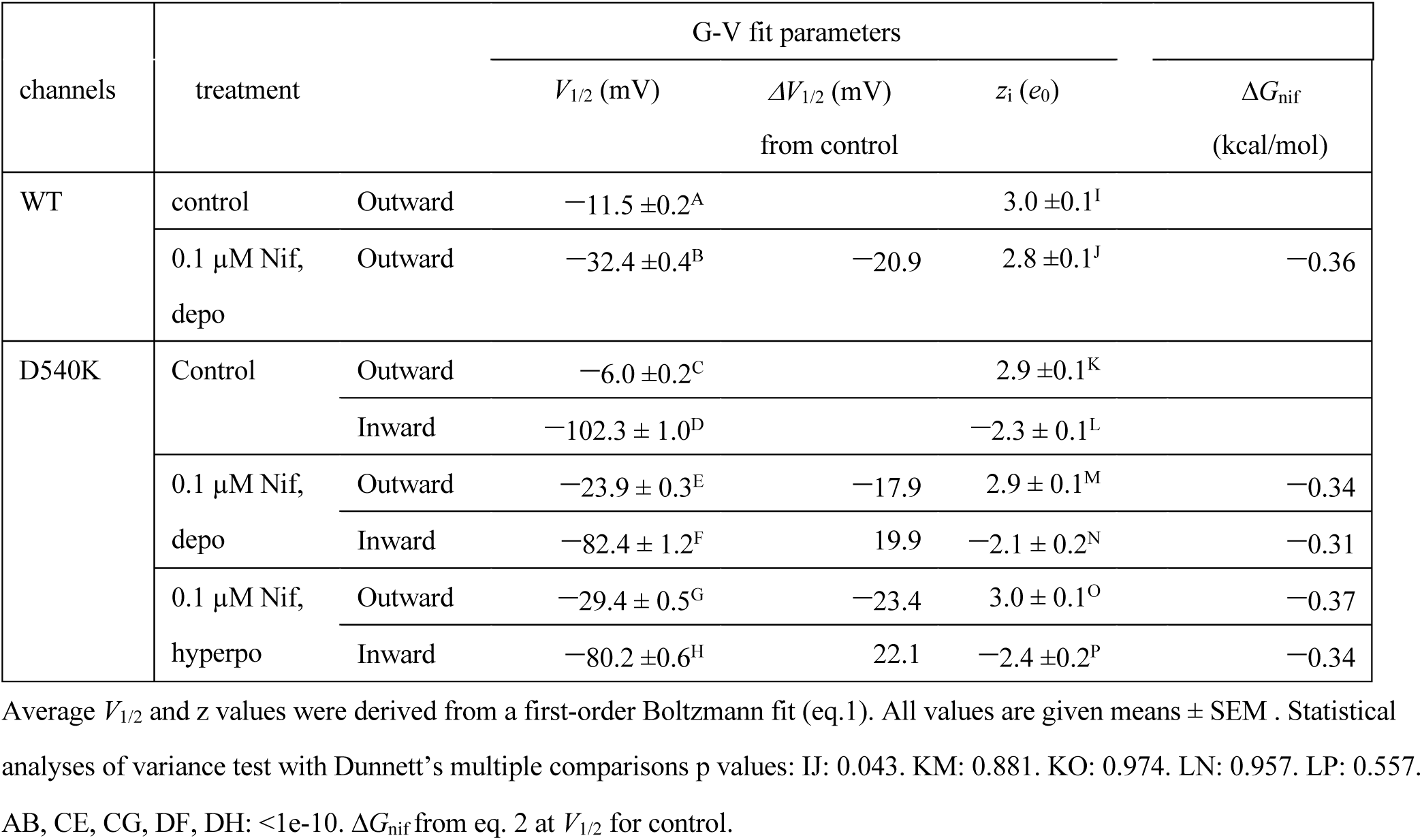
First-order Boltzmann parameter for G-V relationships

## Results

### The facilitation-voltage relationship for nifekalant is similar to the hERG activation-voltage relationship

In *Xenopus* oocytes, the facilitation-voltage relationship for nifekalant is similar to the hERG activation-voltage relationship (Furutani et al., 2019). We assessed whether this was also the case in the HEK293 cells used in this study. We applied nifekalant at 0.1 µM, a concentration which enables facilitation of hERG in HEK293 cells (Furutani et al., 2019). Fig. 1 shows the voltage-dependence of induction of facilitation effect by 0.1 µM nifekalant in WT hERG channels (*V*_1/2_ = −11.4 ± 0.3 mV, *z* = 2.8 ± 0.1 *e*_0_). This voltage dependence is within error of the voltage-dependence of hERG activation (*V*_1/2_ = −11.5 ± 0.2 mV, *z* = 3.0 ± 0.1 *e*_0_), suggesting that a conformational change involved in hERG activation is required for facilitation. Two types of hERG conformational changes might produce this voltage-dependence; outward voltage sensor movement or conformational change in the intracellular gate upon channel activation (Hille, 1977; Hille, 2001). To discriminate between these hypotheses, we altered the coupling between voltage sensor movement and pore opening.

**Fig. 1.**
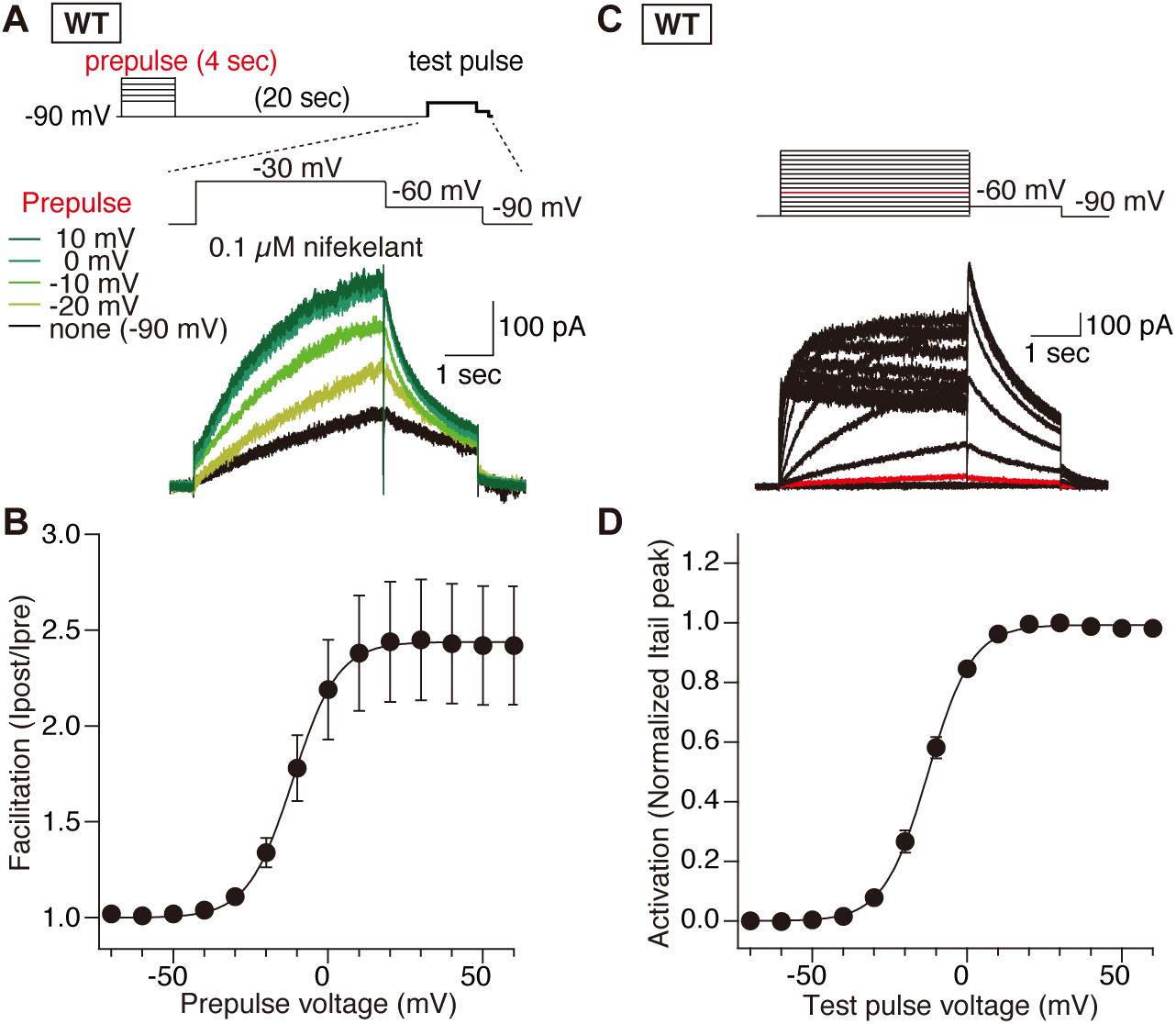
Effects of a preconditioning step depolarization on the induction of facilitation in WT hERG channels. **(A-B)** Effect of a preconditioning step depolarization on the induction of facilitation in WT hERG channels. hERG currents were recorded from HEK293 cells stably expressing hERG at room temperature. 0.1 µM nifekalant was treated. **(A)** The amplitudes of a 4-second preconditioning depolarization (prepulse) were changed from −70 mV to +60 mV by 10 mV increments and the effects on the currents upon the test pulse (−30 mV for 4 sec, then −60 mV for 2 sec) were assayed. Representative traces upon test pulse were shown. **(B)** The relationship between membrane voltage of preconditioning pulse and the steady-state current amplitude. Data are means ± SEM (*n* = 4 cells). The curves were fitted with the Boltzmann function (eq. 1). **(C-D)** Voltage-dependent activation of WT hERG channels. **(C)** The macroscopic hERG currents in response to test pulses step (from −70 to +60 mV by 10 mV increments for 4 sec, then −60 mV for 2 sec) at room temperature; representative traces. **(D)** The relationship between membrane voltage of step pulse and the peak tail current amplitude. Data are means ± SEM (*n* = 6 cells). The curves were fitted with the Boltzmann function (eq. 1).

### Nifekalant is an open-channel blocker of hERG channels and inhibit D540K hERG channels at hyperpolarized potentials

We used D540K hERG to explore the facilitation mechanism because of its unusual activation by hyperpolarization (Mitcheson et al., 2000; Sanguinetti and Xu, 1999). We confirmed the unique gating properties of this D540K hERG. Consistent with the prior reports, current elicited by depolarization from a holding potential of −80 mV appeared to activate instantaneously, followed by rapid inactivation. In the same cell, hyperpolarization from the holding potential of −80 mV activated a small instantaneous current followed by a much slower inward current. The instantaneous component has been proposed to represent channels open at −80 mV D540K hERG (Mitcheson et al., 2000; Sanguinetti and Xu, 1999). We found *V*_1/2_ for depolarization-induced activation = −6.0 ± 0.2 mV, *z* = 2.9 ± 0.1 *e*_0_ and *V*_1/2_ for hyperpolarization-induced activation = −102.3 ± 1.0 mV, *z* = −2.3 ± 0.1 *e*_0_ (Fig. S1).

Before studying the facilitation induction, nifekalant block of D540K hERG channels was evaluated. Nifekalant inhibits WT and D540K hERG channels in a concentration-dependent manner (Fig. 2). The *IC*_50_ of nifekalant inhibition of outward WT hERG current at + 30 mV was 152 ± 8 nM. The *IC*_50_ of D540K hERG at +30 mV was 671 ± 47 nM, indicating a ~4-fold lower inhibitory potency (Fig. 2C). In addition, nifekalant is an even less potent blocker of D540K hERG inward currents, with an *IC*_50_ at −160 mV of 2924 ± 36 nM (Fig. 2B, C). The difference in *IC*_50_ between +30 mV and −160 mV, did not appear to due a charge on nifekalant, as neither the inward nor outward current block had substantial voltage dependence.

**Fig. 2.**
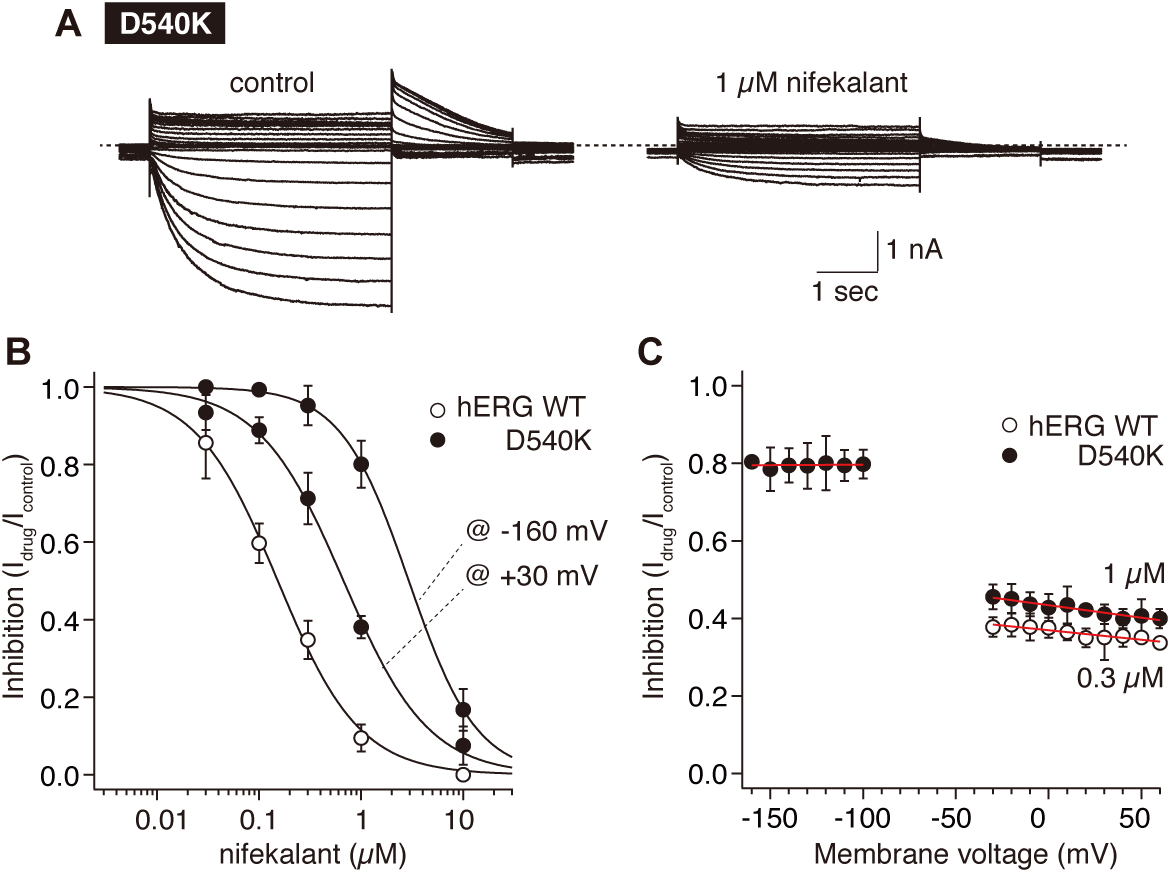
Concentration-dependence block of WT and D540K hERG channels by nifekalant. Effects of nifekalant on WT and D540K hERG channels in HEK293 cells. hERG currents were recorded at room temperature. **(A)** Representative macroscopic D540K hERG currents in response to test pulses step (from −160 to +60 mV by 10 mV increments for 4 sec, then −60 mV for 2 sec). **(B)** Concentration-dependence block of WT and D540K hERG channels by nifekalant. Data are means ± SEM (n = 6-8 cells). The curves were fitted with Hill equation. *IC*_50_ and Hill coefficient (*h*) are 152 ± 8 nM and 1.1 for nifekalant against WT hERG; 671 ± 47 nM and 1.0 against outward D540K hERG currents in HEK293 cells; 2924 ± 36 nM and 1.3 against inward D540K hERG currents in HEK293 cells, respectively. (C) Voltage-dependence block of WT and D540K hERG channels by nifekalant. Concentrations of nifekalant are 0.3 µM and 1 µM against WT and D540K hERG channels, respectively. Data are means ± SEM (n = 4-5 cells). The plots were fitted by a linear.

Some hERG blockers have gated access to their blocking site, where onset of block requires opening of a path to the central cavity. To characterize how inhibition by nifekalant responds to voltage activation of WT and D540K hERG, we designed a 2-pulse protocol containing a brief test-pulse to +10 mV followed by repeating hyperpolarizing pulses to −120 mV which open only D540K but not WT hERG channels (Fig. S1). In 1 µM nifekalant, WT hERG current amplitudes in response to this 2-pulse protocol decay over time (Fig. 3C). Furthermore, no inhibition occurs when cells are held at −80 mV or during repeating hyperpolarizing step-pulses to −120 mV (Fig. 3D), suggesting that hyperpolarization does not gate the path for nifekalant inhibition in WT hERG. In contrast, in 1 µM nifekalant, D540K hERG current progressively declines during repeated hyperpolarizing voltage steps to –120 mV (Fig. 3A, B). As hyperpolarizing pulses to –120 mV open D540K hERG (Fig. S1), these results suggest that opening of the conductance gate allowing nifekalant to access its blocking site, and that outward movement of voltage sensors are not required for access.

**Fig. 3.**
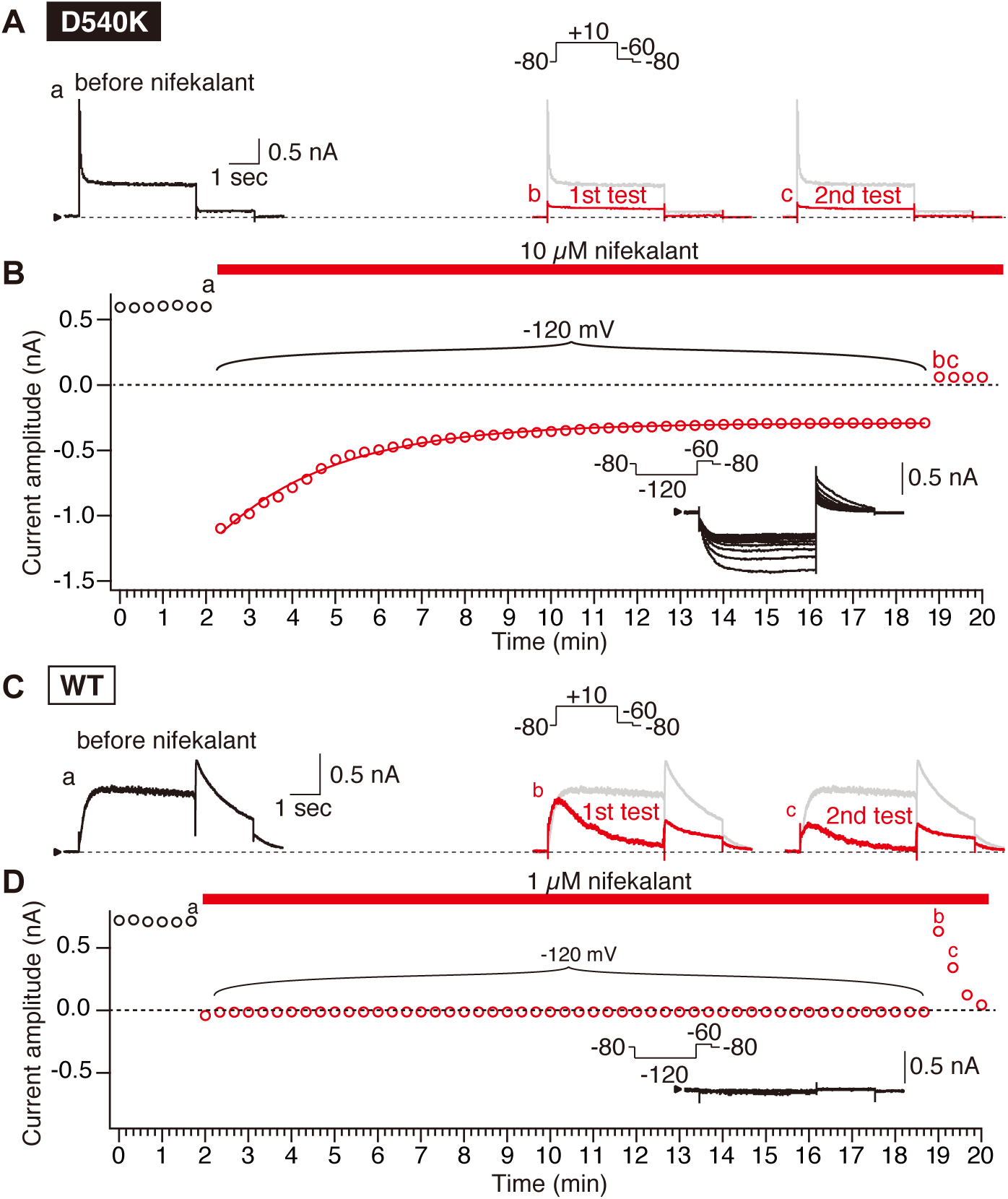
Voltage stimuli are required for nifekalant to inhibit hERG channels. **(A-B)** Effect of hyperpolarizing voltage stimuli for nifekalant to inhibit D540K hERG channels. **(A)** Current traces from a representative cell expressing D540K hERG in vehicle (black and gray) or 10 μM nifekalant (red). Voltage protocol from a holding potential of −80 mV is a 4-second step to +10 mV followed by a 2 sececond-step to −60 mV. Red **a**-**c** indicate time points labeled in **B. (B)** Peak currents during step pulse (circles). Red bar indicates application of 10 μM nifekalant. Insets are representative D540K hERG current traces in response to hyperpolarizing test pulses (−120 mV for 4 sec, then −60 mV for 2 sec) after the treatment of 10 µM nifekalant. Currents evoked by hyperpolarization step to −120 mV in nifekalant are fitted with an exponential function (red curve). Dotted black line indicates Zero current level. **(C-D)** Effect of hyperpolarizing voltage stimuli for nifekalant to inhibit WT hERG channels. **(C)** Current traces from a representative cell expressing WT hERG in vehicle (black and gray) or 1 μM nifekalant (red). Voltage protocol is the same as **A**. Red **a**-**c** indicate time points labeled in **D. (D)** Peak currents during step pulse (circles). Red bar indicates application of 1 μM nifekalant. Insets are representative WT hERG current traces in response to hyperpolarizing test pulses (−120 mV for 4 sec, then −60 mV for 2 sec) after the treatment of 1 µM nifekalant. Dotted black line indicates Zero current level.

### Hyperpolarization-induced currents of D540K hERG can be facilitated by nifekalant

Next, we asked if a depolarizing preconditioning pulse that induces facilitation in WT hERG channels (Furutani et al., 2011; Hosaka et al., 2007; Yamakawa et al., 2012) could induce facilitation in D540K hERG channels. As mentioned above, 0.1 µM nifekalant suppressed the outward currents of D540K hERG activated at depolarized potentials to 88% of control, but not the inward currents activated at depolarized potentials (Fig. 2 and see also Fig. 4CD). In the presence of 0.1 µM nifekalant, we applied a 4-second prepulse to +60 mV, 20-seconds before each test pulse since this voltage protocol is used to monitor the hERG facilitation in WT hERG channels (see Fig. 1). We found that the outward currents of D540K activated at depolarized potentials, especially at potentials close to the threshold, increased after prepulse to +60 mV in 0.1 µM nifekalant (Fig. 4C, D). The *V*_1/2_ of activation of the facilitated fraction was −23.9 ± 0.3 mV, shifted negative from the control (Δ*V*_1/2_ = −17.9 mV), by roughly the same degree as WT (Δ1*V*_1/2_ = −20.9 mV)(Furutani et al., 2019). Such a shift is a hallmark feature of facilitation by nifekalant and other facilitating blockers (Carmeliet, 1993; Furutani et al., 2019; Furutani et al., 2011; Hosaka et al., 2007; Jiang et al., 1999; Perry et al., 2010; Yamakawa et al., 2012).

**Fig. 4.**
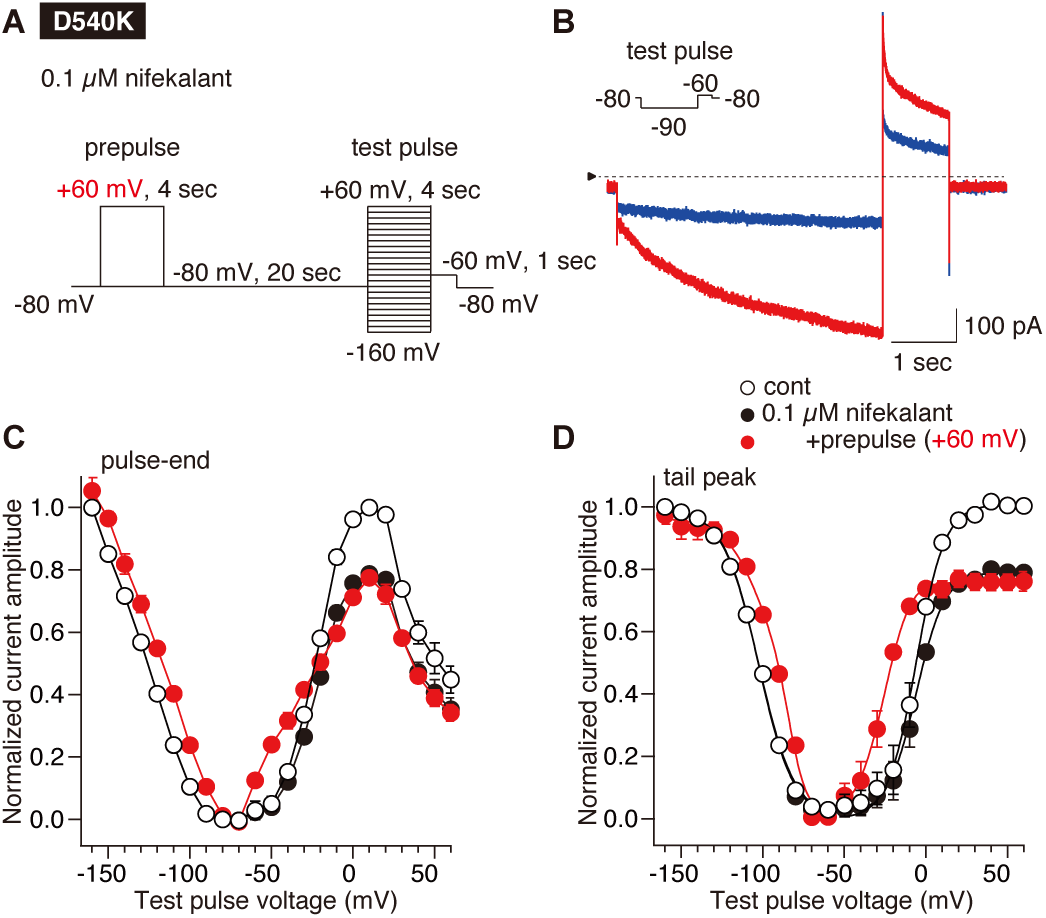
Nifekalant facilitates D540K hERG activation at both depolarized and hyperpolarized potentials. **(A)** Pulse protocol of this experiment. The amplitudes of a 4-second preconditioning depolarization (prepulse) were +60 mV and the effects on the currents from HEK cells expressing D540K hERG upon the test pulse (from −160 mV to +60 mV by 10 mV increments for 4 sec, then −60 mV for 2 sec) were assayed. **(B)** Superimposed D540K hERG currents recorded in the same cell before (cont, black) and after perfusion of 0.1 μM nifekalant with (+prepulse, red) or without (blue) the preconditioning pulse of +60 mV. **(C-D)** The relationship between membrane voltage of test pulse and the steady-state current amplitude **(C)** and the tail current amplitude (activation curve) **(D). (C)** The amplitudes of hERG step currents in the absence (open circles), and in the presence of 0.1 μM nifekalant with (filled black circles) or without (filled red squares) the preconditioning pulse were measured at the pulse-end during the test pulse to indicated voltages. Outward currents recorded at −50 mV to +60 mV was normalized to the currents evoked by voltage steps to +10 mV in the absence of nifekalant. Inward currents recorded at −160 mV to −60 mV was normalized to the currents evoked by voltage steps to −160 mV in the absence of nifekalant. **(D)** Voltage-dependent hERG activation. The amplitudes of hERG tail currents in the absence (open circles), and in the presence of 0.1 μM nifekalant with (filled black circles) or without (filled red squares) the preconditioning pulse were measured at the peak after the test pulse to indicated voltages. Depolarization-activated tail currents recorded at −60 mV was normalized to the currents evoked by voltage steps to +10 mV in the absence of nifekalant. Hyperpolarization-activated tail currents recorded at −60 mV was normalized to the currents evoked by voltage steps to −160 mV in the absence of nifekalant. Data are means ± SEM (n = 5 cells). In activation curve **(D)** the curves were fitted with the single (for control and nifekalant without prepulse) or double Boltzmann equation (for nifekalant with prepulse).

Interestingly, prepulse to +60 mV in 0.1 µM nifekalant also increases the inward currents of D540K (Fig. 4 B-D). The voltage-midpoint of the hyperpolarization-activated D540K currents was −82.4 ± 1.2 mV, shifted to more positive voltages by 19.9 mV. This interesting result reveals that nifekalant can facilitate both the usual depolarization activation mechanism and the unusual hyperpolarization activation mechanism of D540K hERG.

### Hyperpolarization can induce hERG facilitation in D540K, but not WT, hERG

The hypothesis that facilitation is induced by stimuli that activate the hERG conductance predicts that facilitation will be also induced when the D540K hERG conductance is activated by hyperpolarization. To examine this possibility, we subjected D540K cells to a hyperpolarizing prepulse in the presence of nifekalant. First, we experimented with 0.1 µM nifekalant, the concentration in Fig. 4. We found that a 4-second hyperpolarizing prepulse to −160 mV could not induce facilitation (Fig. 5). We reasoned this could be because the 0.1 µM concentration of nifekalant is dilute to interact sufficiently with D540 hERG channels at −160 mV, as 0.1 µM does not appreciably block inward currents activated at hyperpolarized potentials (Fig. 2), and facilitation occurs only with concentrations that also block channels (Furutani et al., 2011). To block D540K hERG at –160 mV to a similar degree as 0.1 µM nifekalant blocked at +20 mV (19%, see Fig. 4CD), we increased nifekalant to 1 µM to block inward current of D540K hERG channels at −160 mV by 20%. In 1 µM nifekalant, a –160 mV prepulse induced facilitation (Fig. 6). The facilitated *V*_1/2_ for the currents activated at depolarized potentials and hyperpolarized potentials were −29.4 ± 0.5 mV (Δ*V*_1/2_ from the unfacilitated *V*_1/2_ = −23.4 mV) and −80.2 ± 0.6 mV (Δ*V*_1/2_ = 22.1 mV), respectively. Thus, in 1 µM nifekalant, a –160 mV prepulse induced facilitation similar to a +60 mV prepulse in 0.1 µM.

**Fig 5.**
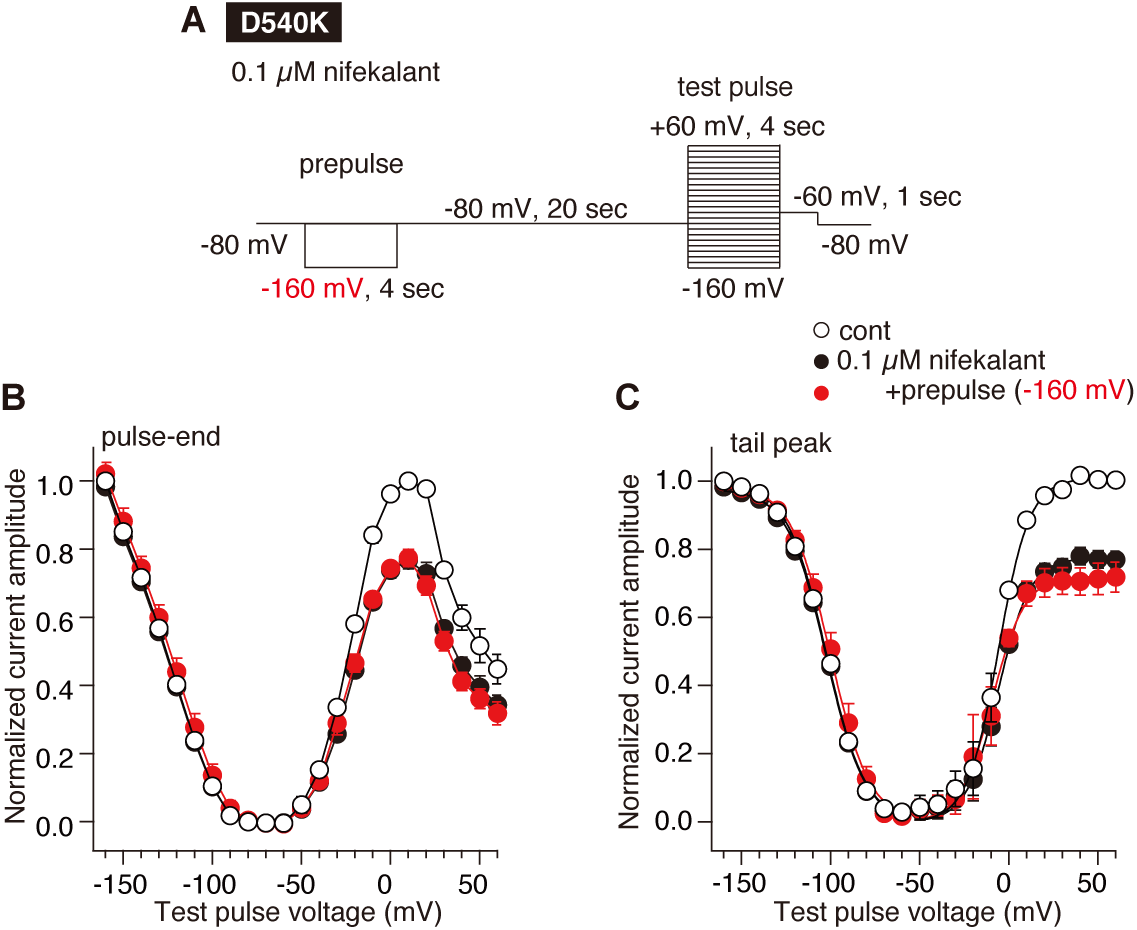
Hyperpolarizing prepulse does not induce facilitation effect on D540K hERG activation in the presence of 0.1 µM of nifekalant. **(A)** Pulse protocol of this experiment. The amplitudes of a 4-second preconditioning hyperpolarization (prepulse) were −160 mV and the effects on the currents from HEK cells expressing D540K hERG upon the test pulse (from −160 mV to +60 mV by 10 mV increments for 4 sec, then −60 mV for 2 sec) were assayed. **(B-C)** The relationship between membrane voltage of test pulse and the steady-state current amplitude **(B)** and the tail current amplitude (activation curve) **(C). (B)** The amplitudes of hERG step currents in the absence (open circles), and in the presence of 0.1 μM nifekalant with (filled black circles) or without (filled red squares) the preconditioning pulse were measured at the pulse-end during the test pulse to indicated voltages. Outward currents recorded at −50 mV to +60 mV was normalized to the currents evoked by voltage steps to +10 mV in the absence of nifekalant. Inward currents recorded at −160 mV to −60 mV was normalized to the currents evoked by voltage steps to −160 mV in the absence of nifekalant. **(C)** Voltage-dependent hERG activation. The amplitudes of hERG tail currents in the absence (open circles), and in the presence of 0.1 μM nifekalant with (filled black circles) or without (filled red squares) the preconditioning pulse were measured at the peak after the test pulse to indicated voltages. Depolarization-activated tail currents recorded at −60 mV was normalized to the currents evoked by voltage steps to +10 mV in the absence of nifekalant. Hyperpolarization-activated tail currents recorded at −60 mV was normalized to the currents evoked by voltage steps to −160 mV in the absence of nifekalant. Data are means ± SEM (n = 5 cells). In activation curve **(C)** the curves were fitted with the single Boltzmann function (eq. 1).

**Fig 6.**
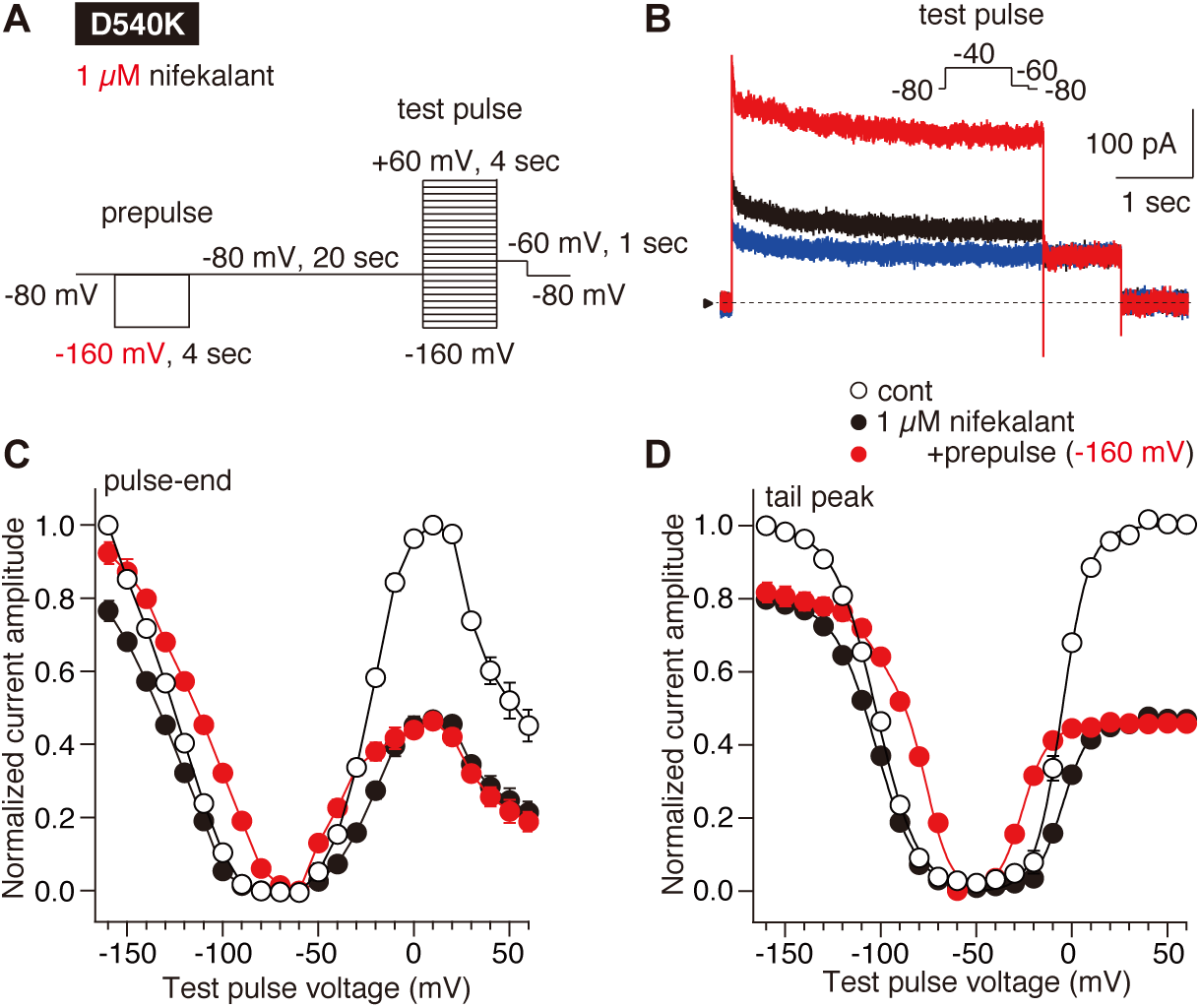
Hyperpolarizing prepulse induces facilitation effect on D540K hERG activation in the presence of 1 µM of nifekalant. **(A)** Pulse protocol of this experiment. The amplitudes of 4-second preconditioning hyperpolarization (prepulse) were −160 mV and the effects on the currents from HEK cells expressing D540K hERG upon the test pulse (from −160 mV to +60 mV by 10 mV increments for 4 sec, then −60 mV for 2 sec) were assayed. **(B)** Superimposed hERG currents recorded in the same cell before (cont, black) and after perfusion of 1 μM nifekalant with (+prepulse, red) or without (blue) the preconditioning pulse of −160 mV. **(C-D)** The relationship between membrane voltage of test pulse and the steady-state current amplitude **(C)** and the tail current amplitude (activation curve) **(D). (C)** The amplitudes of hERG step currents in the absence (open circles), and in the presence of 0.1 μM nifekalant with (filled black circles) or without (filled red squares) the preconditioning pulse were measured at the pulse-end during the test pulse to indicated voltages. Outward currents recorded at −50 mV to +60 mV was normalized to the currents evoked by voltage steps to +10 mV in the absence of nifekalant. Inward currents recorded at −160 mV to −60 mV was normalized to the currents evoked by voltage steps to −160 mV in the absence of nifekalant. **(D)** Voltage-dependent hERG activation. The amplitudes of hERG tail currents in the absence (open circles), and in the presence of 1 μM nifekalant with (filled black circles) or without (filled red squares) the preconditioning pulse were measured at the peak after the test pulse to indicated voltages. Depolarization-activated tail currents recorded at −60 mV was normalized to the currents evoked by voltage steps to +10 mV in the absence of nifekalant. Hyperpolarization-activated tail currents recorded at −60 mV was normalized to the currents evoked by voltage steps to −160 mV in the absence of nifekalant. Data are means ± SEM (n = 5 cells). In activation curve **(D)** the curves were fitted with the single (for control and nifekalant without prepulse) or double Boltzmann equation (for nifekalant with prepulse).

To eliminate the possibility that hyperpolarizing prepulses induce facilitation of WT hERG, we conducted controls with the same hyperpolarizing prepulse protocol on WT hERG. Facilitation was not induced by a –160 mV prepulse with either 0.1 µM or 1 µM nifekalant (Fig. 7).

**Fig. 7.**
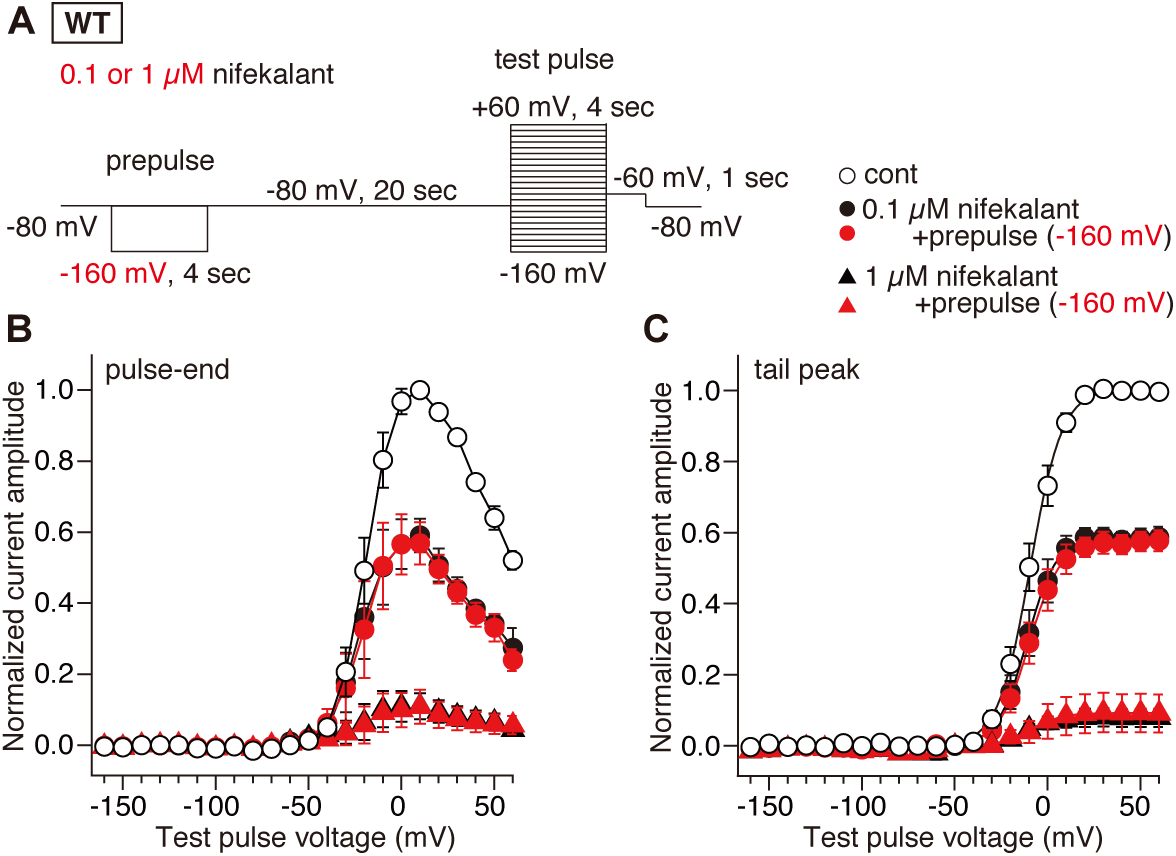
Hyperpolarizing prepulse does not induce facilitation effect on WT hERG activation in the presence of 0.1 or 1 µM of nifekalant. **(A)** Pulse protocol of this experiment. The amplitudes of a preceding hyperpolarization (prepulse) were −160 mV and the effects on the currents from HEK cells expressing WT hERG upon the test pulse (from −160 mV to +60 mV by 10 mV increments for 4 sec, then −60 mV for 2 sec) were assayed. **(B-C)** The relationship between membrane voltage of test pulse and the steady-state current amplitude **(B)** and the tail current amplitude (activation curve) **(C). (B)** The amplitudes of hERG step currents in the absence (open circles), and in the presence of 0.1 μM nifekalant with (filled black circles) or without (filled red circles), or in the presence of 1 μM nifekalant with (filled black triangles) or without (filled red triangles) the preconditioning pulse were measured at the pulse-end during the test pulse to indicated voltages. Outward currents recorded at −50 mV to +60 mV was normalized to the currents evoked by voltage steps to +10 mV in the absence of nifekalant. Inward currents recorded at −160 mV to −60 mV was normalized to the currents evoked by voltage steps to −160 mV in the absence of nifekalant. **(C)** Voltage-dependent hERG activation. The amplitudes of hERG tail currents in the absence (open circles), and in the presence of 0.1 μM nifekalant with (filled black circles) or without (filled red circles), or in the presence of 1 μM nifekalant with (filled black triangles) or without (filled red triangles) the preconditioning pulse were measured at the peak after the test pulse to indicated voltages. Depolarization-activated tail currents recorded at −60 mV was normalized to the currents evoked by voltage steps to +10 mV in the absence of nifekalant. Hyperpolarization-activated tail currents recorded at −60 mV was normalized to the currents evoked by voltage steps to −160 mV in the absence of nifekalant. Data are means ± SEM (n = 5-8 cells). In activation curve **(C)** the curves were fitted with the single Boltzmann function (eq. 1).

These results indicate that to induce facilitation by nifekalant: 1) a preconditioning pulse that activates hERG is required, and 2) nifekalant must block hERG during the preconditioning pulse.

## Discussion

These results with D540K hERG advance our understanding of the mechanism of hERG channel facilitation by blocking agents. Specifically, these results clarify that facilitation is a stabilization of the open conformation of the pore rather than a negative shift of voltage sensitivity, and that the prepulse which induces facilitation acts by opening a gate within the channel.

### What is the mechanism underlying the induction of facilitation by a prepulse?

A preconditioning pulse to a depolarized or positive potential is required to induce facilitation by nifekalant (Furutani et al., 2011; Hosaka et al., 2007). This depolarization-dependence of the induction of facilitation is a common characteristic of facilitation induction for other hERG channel inhibitors (Carmeliet, 1993; Furutani et al., 2011; Jiang et al., 1999; Perry et al., 2010; Yamakawa et al., 2012). In this study, we find that activation of the hERG channel conductance, not depolarization *per se*, is required to induce facilitation, as activating the D540K conductance by hyperpolarization also induces facilitation. Why is activation of the hERG conductance required to induce facilitation? We propose that activation of the hERG conductance allows the hERG blocker to access a facilitation site. Voltage-gated ion channel activation involves opening an S6 intracellular gate which can allow channel blockers to access a binding site in the central cavity of channel (Asai et al., 2021; del Camino and Yellen, 2001; Hosaka et al., 2007; Sanguinetti and Tristani-Firouzi, 2006; Vandenberg et al., 2012). Alanine-scanning mutagenesis has suggested that amino acid residues of the central cavity around the pore helix, such as T623, S624, V625 and on S6 helix, such as G648, Y652 and F656, are important for facilitation (Hosaka et al., 2007). Thus, it is likely that facilitating hERG blockers enter the central cavity when the intracellular gate is opened. This idea is supported by the finding that nifekalant must block during a prepulse to induce facilitation, whether the prepulse activates D540K by depolarization or hyperpolarization (Fig. 4, 5). This suggests that when facilitation is induced, the membrane potential must open the intracellular gate such that the drug can enter a site in the central cavity. It is unclear if the drug binds similarly to block and facilitate. A previous mutagenesis study showing different impacts of single amino substitutions on block and facilitation suggests that the molecular interactions for facilitation are different than for block (Hosaka et al., 2007). Still, both interactions occur in the central cavity. Therefore, activation of the voltage sensor may open the path to the central cavity, to both block and facilitation.

### How does nifekalant to affect activation of hERG?

How nifekalant facilitates activation remains a question. The present study demonstrates that nifekalant can also facilitate hyperpolarization-induced activation of D540K hERG channels. Thus, the polarity of the facilitation effect is malleable, indicating that facilitation does not simply stabilize of voltage sensors in an up state. This result suggests that nifekalant acts on a shared mechanism underlying D540K hERG openings by depolarization and hyperpolarization. Although the mechanism by which D540K hERG opens at hyperpolarized potentials is not completely clear, it involves a change in coupling between voltage-sensor and pore domains leading to hyperpolarization-triggered opening of the intracellular S6 gate (Tristani-Firouzi et al., 2002). As the facilitation effect tracks conductance, this suggests that facilitation responds to and alters pore dynamics directly, rather than acting through voltage sensors.

Blockers which bind within the central cavity of Kv channels can alter channel gating. For example, the pore blocker 4-aminopyridine requires opening of the S6 gate of Shaker Kv1 channels to block, and then stabilizes pores in closed conformations (Armstrong and Loboda, 2001). Interestingly, 4-aminopyridine can cause an increase in the conductance of Kv6.4/Kv2.1 heteromeric channels (Stas et al., 2015). The Kv2 blocker RY785 accelerates channel deactivation to trap itself (Marquis and Sack, 2022). On the other hand, quaternary ammonium ions such as tetraethylammonium stabilize open channels, albeit in a blocked conformation and this can cause a shift of the conductance voltage relation to more negative voltages (Armstrong, 1971; Holmgren et al., 1997; Melishchuk and Armstrong, 2001). We suggest that stabilization of an open S6 gate by blockers could play a role in facilitation.

### Nifekalant block varies with current direction, not membrane voltage

The nifekalant block of D540K hERG inward currents was weaker than block of outward currents (Fig. 2). Nifekalant has pKa of 7.05, suggesting 30% of nifekalant is positively charged in the recording solution at pH 7.4. However, little voltage dependence can be attributed to charge on nifekalant, as the fractional block of either inward or outward K^+^ currents changed little with voltage (Fig. 2C). Thus, the difference between inward and outward currents could arise from an interaction of permeant K^+^ ions with a drug trapped within the pore (Pareja et al., 2013), a difference in inactivation, or another conformational change of the channel. Mutation of S4-S5 linker has only minor effects on the rate of hERG channel inactivation, and D540K hERG channels are inactivated at depolarized potentials and less inactivated at hyperpolarized potentials (Sanguinetti and Xu, 1999). Nifekalant may have a higher affinity to the inactivated state than the activated state (Kushida et al., 2002), and this could also possibly account for the weaker affinity, as the inward currents of D540K do not inactivate. In addition, nifekalant has a weaker efficacy against D540K than WT. This is the also the case for hERG blockers such as MK-499 (Mitcheson et al., 2000), and the mechanism undergirding this efficay difference between WT vs. D540K is not well understood. Previous studies with D540K hERG do not rule out the possibility that reduced binding affinity to the mutant channels results from an allosteric effect unrelated to channel state.

### Insights into the risk of cardiac arrhythmias

hERG blockers carry a risk of lethal arrhythmias. Mechanism-based quantitative prediction of hERG-associated drug-induced arrhythmias has been a challenge (Gintant et al., 2016; Li et al., 2020; Sager et al., 2014). The present finding that channel activation is essential for facilitation suggests a link between blockade and facilitation, which may be help assess the arrhythmic risk of drugs. We previously conducted simulations suggesting that facilitation by hERG blockers may decrease incidence of early afterdepolarizations in cardiac myocytes (Furutani et al., 2019). In that study, we used experimental data with nifekalant as the representative drug to block and facilitate hERG channels. Because the exact mechanism of induction of and recovery from facilitation was unknown, we modeled facilitation based on experimentally obtained dose-responses curves without a mechanistic underpinning. Further clarification of the mechanism of the facilitation effect could enable for a more refine theoretical approach for prediction of arrhythmia risk.

## Conclusion

In this study, we analyzed the facilitation effect of nifekalant on hyperpolarization-evoked D540K hERG channel activation and demonstrated that nifekalant exerts a facilitation effect on hERG currents at hyperpolarized potentials as well as depolarized potentials. The facilitation effect is induced by the drug interaction with hERG channels while their conductance is activated.

## Abbreviations

HEK: human embryonic kidney
hERG: human ether-à-go-go-related gene
*I*_Kr_: rapid component of the delayed-rectifier potassium current
TdP: *Torsade de Pointes*
*V*_1/2_: half-activation voltage
WT: wild-type

## Acknowledgments

We thank Drs. Yoshihisa Kurachi, Atsushi Inanobe, Hiroshi Hibino (Osaka University) and Mitsuhiro Yamada (Shinsyu University) for their support to set up facilities and equipment at Department of Pharmacology in Tokushima Bunri University. We are grateful to Dr. Mark T. Keating and Dr. Michael C. Sanguinetti (University of Utah) for providing us with hERG clone, Dr. Craig T. January (University of Wisconsin) for providing us with HEK293 cell lines stably expressing hERG.

## Authorship Contributions

Participated in research design: Furutani, Sack, and Kita

Conducted Experiments: Furutani, Kawano, Ichiwara, and Adachi

Contributed new reagents or analytic tools: Furutani, Adachi, Clancy, and Sack

Performed data analysis: Furutani and Sack

Wrote or contributed to the writing of the manuscript: The manuscript was drafted by Furutani,

Sack, and Kita and was revised and corrected by all co-authors.

## Conflict of Interest statement

The authors declare no competing financial interests.

## Figure Legends

**Fig S1.**
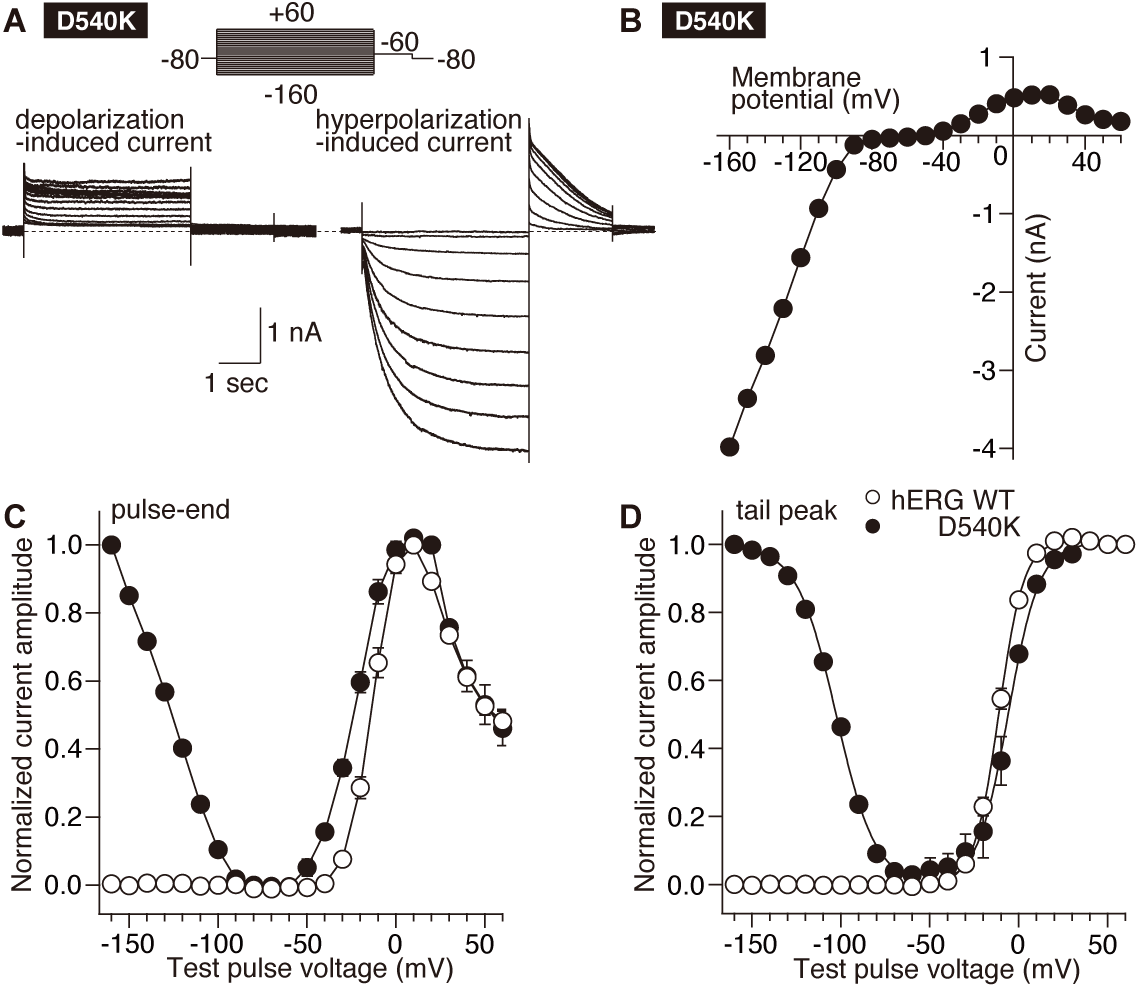
Electrophysiological properties of D540K hERG channels expressed in HEK293 cells. **(A)** Representative whole cell recordings of D540K hERG currents elicited by 4-sec depolarizing test pulses from −80 to +60 mV in 10-mV increments **(left)** and by 4-sec hyperpolarized test pulses from −80 to −160 mV in 10-mV decrements **(right). (B)** Representative steady state I-V relationship for D540K hERG. **(C-D)** The relationship between membrane voltage of test pulse and the steady-state current amplitude **(C)** and the tail current amplitude (activation curve) **(D). (C)** The amplitudes of hERG step currents in WT (open circles) and D540K hERG (filled black circles) were measured at the pulse-end during the test pulse to indicated voltages. Outward currents recorded at −50 mV to +60 mV was normalized to the currents evoked by voltage steps to +10 mV. Inward currents recorded at −160 mV to −60 mV was normalized to the currents evoked by voltage steps to −160 mV. **(D)** Voltage-dependent hERG activation. The amplitudes of hERG tail currents in WT (open circles) and D540K hERG (filled black circles) were measured at the peak after the test pulse to indicated voltages. Depolarization-activated tail currents recorded at −60 mV was normalized to the currents evoked by voltage steps to +10 mV. Hyperpolarization-activated tail currents recorded at −60 mV was normalized to the currents evoked by voltage steps to −160 mV. Data are means ± SEM (n = 5 cells). In activation curve **(D)**, the curves were fitted with the single Boltzmann function (eq. 1).

## Notes

This study was supported by the Scientific Research (C) 21K06812 (K.F.) from the Ministry of Education, Science, Sports and Culture of Japan, the Japan Society for the Promotion of Science, and NIH grants U01HL126273 and R01HL128537 (K.F. C.E.C. and J.T.S.).

### Competing Interest Statement

The authors have declared no competing interest.

